# Growth of soil ammonia-oxidizing archaea on air-exposed solid surface

**DOI:** 10.1101/2024.07.15.603479

**Authors:** Christiana Abiola, Joo-Han Gwak, Ui-Ju Lee, Samuel Imisi Awala, Man-Young Jung, Woojun Park, Sung-Keun Rhee

## Abstract

Soil microorganisms often thrive as microcolonies or biofilms within pores of soil aggregates exposed to the soil atmosphere. However, previous studies on the physiology of soil ammonia-oxidizing microorganisms (AOM), which play a critical role in the nitrogen cycle, were primarily conducted using freely suspended AOM cells (planktonic cells) in liquid media. In this study, we examined the growth of two representative soil ammonia-oxidizing archaea (AOA), *Nitrososphaera viennensis* EN76 and “*Nitrosotenuis chungbukensis*” MY2, and an ammonia-oxidizing bacterium, *Nitrosomonas europaea* ATCC 19718 on polycarbonate membrane filters floated on liquid media to observe their adaptation to air-exposed solid surfaces. Interestingly, ammonia oxidation activities of *N. viennensis* EN76 and “*N. chungbukensis*” MY2 were significantly repressed on floating filters compared to the freely suspended cells in liquid media. Conversely, the ammonia oxidation activity of *N. europaea* ATCC 19718 was comparable on floating filters and liquid media. *N. viennensis* EN76 and *N. europaea* ATCC 19718 developed microcolonies on floating filters. Transcriptome analysis of *N. viennensis* EN76 floating filter-grown cells revealed upregulation of unique sets of genes for cell wall and extracellular polymeric substance biosynthesis, H_2_O_2_-induced oxidative stress defense, and ammonia oxidation, including ammonia monooxygenase subunit C (*amoC3*) and the multicopper oxidases. These genes may play a pivotal role in adapting AOA to air-exposed solid surfaces. Furthermore, the floating filter technique resulted in the enrichment of distinct soil AOA communities dominated by the “*Ca*. Nitrosocosmicus” clade. Overall, this study sheds light on distinct adaptive mechanisms governing AOA growth on air-exposed solid surfaces.

## Introduction

Ammonia-oxidizing microorganisms (AOM) play a crucial role in the global nitrogen cycle in terrestrial environments [1, 2]. They include ammonia-oxidizing Archaea (AOA) and ammonia-oxidizing bacteria (AOB), which mediate the first and rate-limiting step of nitrification: ammonia (NH_3_) oxidation to nitrite (NO_2_^-^) [3, 4]. Another group of bacterial ammonia-oxidizers, known as complete ammonia-oxidizers (comammox), can oxidize NH_3_ to nitrate (NO_3_^-^) via NO_2_^-^ in terrestrial environments [5, 6].

Terrestrial AOA constitute a significant portion of the soil nitrification microbiome [1, 7]. AOA belong to the class *Nitrososphaeria*, which is affiliated with the phylum *Nitrososphaerota*, formerly known as *Thaumarchaeota* [8, 9]. They can be classified into four major orders: “*Candidatus* Nitrosocaldales”*, Nitrosopumilales,* “*Ca.* Nitrosotaleales,” and *Nitrososphaerales* [10]. Recently, Zheng *et al*. [11] reported “*Ca.* Nitrosomirales”, a novel order of the AOA widespread in terrestrial and marine environments. *Nitrososphaerales* is mainly a group of soil-dwelling AOA, also known as group I.1b [12]. They have some cultivated representatives and various subclusters that have not been cultured [13–15]. The isolated strains of *Nitrososphaerales* are obligate aerobes, mesophiles/moderately thermophiles with chemolithoautotrophic metabolism through ammonia oxidation and CO_2_ fixation [16–18].

Given that most soil microorganisms proliferate by colonizing mineral surfaces [19], which are often directly exposed to the soil atmosphere within pores of soil aggregates. Surprisingly, the primary method for cultivating soil AOM remains the liquid culture system. Previous nitrification studies have mainly focused on strains isolated from liquid culture systems, and their physiology has been studied exclusively in liquid media. It is worth noting that AOM strains obtained through the liquid culture system may exhibit notable variations in their physiology when cultivated on a solid interface. These variations may arise from differences in substrate accessibility and chemical disparities at the solid-liquid interface. Hence, fundamental questions about their physiology within the pores of soil aggregates still need to be explored. To address this gap, we adopted the floating filter cultivation technique, previously described for enriching and isolating ubiquitous uncultivated bacteria [20–22], to simulate solid systems and investigate AOA adaptation to air-exposed solid surfaces. Although the floating filter cultivation technique has not been reported for AOA, a few studies have described the formation of microcolonies on membrane filters by AOB strains belonging to the genera *Nitrosomonas* and *Nitrosospira* [23, 24]. However, detailed information about their physiology when grown on these solid surfaces remains elusive.

Notably, the soil *Nitrososphaerales* typically grows in aggregates, implying the ability to extensively modify their cell to adhere to surfaces and form biofilm [12, 16, 18]. Thus, we compare the growth and physiological properties of soil-dwelling AOA from two phylogenetically distinct clades, *Nitrosopumilales* and *Nitrososphaerales*, along with an ammonia-oxidizing bacterium from the genus *Nitrosomonas*, on floating membrane filters and liquid media. We demonstrated that the physiology of AOA grown on air-exposed solid surfaces was different from those grown as freely suspended cells in liquid media. Using transcriptomic analysis, we revealed the distinct response mechanisms of these strains to the floating filter. Finally, the floating filter cultivation technique was demonstrated to enrich distinct soil surface-adapted AOA communities from agricultural soil.

## Materials and methods

### Cultivation in the liquid media

For this study, we selected representative strains of soil AOA of *Nitrosopumilales* and *Nitrososphaerales* clades, □*Nitrosotenuis chungbukensis*” MY2 and *Nitrososphaera viennensis* EN76, respectively, together with an ammonia-oxidizing bacterium, *N. europaea* ATCC 19718. These pure AOM strains were cultivated in artificial freshwater medium (AFM) under optimal conditions as previously described [25–27]. Briefly, pure cultures of *N. viennensis* EN76, □*N. chungbukensis*” MY2, and *N. europaea* ATCC 19718 were incubated in the dark without shaking at their optimum growth temperature and pH. The medium pH was 7.5 for the AOA strains and 7.8 for *N. europaea* ATCC 19718. Ammonium chloride (NH_4_Cl) (1 mM) from pre-sterilized stocks was added as the sole energy source. The growth medium of *N. viennensis* EN76 and □*N. chungbukensis*” MY2 was always supplemented with sodium pyruvate (0.1 mM) as an H_2_O_2_ scavenger [25, 28]. The pH of the media was adjusted with sterile NaOH when necessary. The growth and ammonia oxidation activities of all strains were monitored by nitrite (NO_2_^-^) accumulation. NO_2_^-^ concentration was measured colorimetrically using the Griess test as previously described [26]. Cultures were regularly monitored for contamination of heterotrophic bacteria by checking turbidity in Luria-Bertani and Reasoner’s 2A broth (1:10).

### Cultivation on the floating filters

We compared the growth and ammonia oxidation activities of three AOM strains, *N. viennensis* EN76, □*N. chungbukensis*” MY2, and *N. europaea* ATCC 19718 cells on floating filters and the control culture grown as freely suspended cells in AFM (hereafter referred to as control culture in liquid media). *N. europaea* ATCC 19718, *N. viennensis* EN76, and □*N. chungbukensis*” MY2, which had estimated cell densities of ∼ 10^6^ cells mL^-1^, 10^7^ cells mL^-1^, and 10^8^ cells mL^-1^, respectively, after oxidizing 1mM ammonia, were serially diluted in 10-fold increments. A 1-mL aliquot of each dilution was used as an inoculum for floating filters and control culture in liquid media. Inoculum size hereafter is expressed as the number of cells in the 1-mL aliquot of each dilution. For the floating filter, pre-rinsed sterile nucleopore polycarbonate (PC) track-etched filters (Cytiva, Marlborough, MA, US) with a diameter of 47 mm and a pore size of 0.1 µm were used for the experiment. The filter was mounted on the autoclaved filter holders. Afterward, the 1 mL inoculum was mixed with 4 mL AFM and vacuum-filtered with < 10 mm Hg. Then, the filter was aseptically placed on top of 10 mL AFM in a Petri dish using sterile tweezers. The Petri dishes were placed in a sealed container and incubated in the dark without shaking at 42 °C, 30 °C, and 25 °C for *N. viennensis* EN76, □*N. chungbukensis*” MY2, and *N. europaea* ATCC 19718, respectively. A blank filter (uninoculated filter) was included as a negative control. The ammonia oxidation activities of all strains on floating filters were monitored by measuring NO_2_^-^ accumulation in AFM below the filter. Growth rates were calculated based on the assumption that NO_2_^-^ production is correlated to the growth of ammonia oxidizers [29]. For biomass determination, cells of microcolonies were washed twice from the filter and harvested using centrifugation (13000 x *g*, 7min, 25 °C). The total cellular protein was quantified to determine biomass production using the micro BCA protein assay kit (Thermo Fisher Scientific, Waltham, MA, USA), according to the manufacturer’s instructions.

### Microscopy

A fluorescence microscope was used to monitor the growth of AOM in microcolonies on the floating filter. The counterstain DAPI (blue) was added to reveal the size, morphology, and arrangement of the microcolonies of AOM on the filter. For staining, about 30 µl of 200 µg mL^−1^ DAPI (4,6-diamidino-2-phenylindole) was added on top of filters (cells side up), after which filters were incubated for 10 mins and dried at 37 °C in the dark. The dried filters were mounted on glass slides with coverslip and viewed under 1000x with immersion oil. Microcolonies and cells were imaged using an Olympus BX61 fluorescence microscope with a U-MWU2 fluorescence mirror unit.

### Transcriptomic analysis

Transcriptome analysis was performed on cells of *N. viennensis* EN76 (triplicates) and *N. europaea* ATCC 19718 (> triplicates) grown on floating filters and the control culture in liquid media. At first, a 10-mL AFM was used to cultivate cells of *N. viennensis* EN76 and *N. europaea* ATCC 19718 on floating filters (in Petri dishes) and the control culture in liquid media (in cell culture flasks) under optimum conditions. Before the end of ammonia oxidation, once the NO_2_^-^ had accumulated to ∼ 0.7 mM, a further 1 mM NH_4_Cl was added to sustain exponential growth, and the cultures were scaled up to a vessel containing 300 mL AFM to obtain sufficient biomass. To scale up the volume of AFM for the floating filter-grown cells, the filters were transferred from Petri dishes to bottles containing 300 mL fresh AFM with 1 mM NH_4_Cl. The total RNA was extracted from cells of the scaled-up filters, and the control cultures in liquid media (harvested by filtration) after ∼ 0.6 mM NH_3_ was oxidized. RNA extraction was performed using the AllPrep DNA/RNA Mini Kit (Qiagen) following the manufacturer’s instructions.

RNA quality check was performed with the Agilent 2100 Expert Bioanalyzer (Agilent), and cDNA libraries from the RNA samples were prepared using the Nugen Universal Prokaryotic RNA-Seq Library Preparation Kit. The cDNA libraries were sequenced via NovaSeq6000 (Illumina) at LabGenomics (Seongnam, Korea). Reads quality was assessed by FastQC (v0.11.8) [30]. For trimming reads, Trimmomatic (v0.36) [31] was used with the options: SLIDINGWINDOW:4:15 LEADING:3 TRAILING:3 MINLEN:38 HEADCROP:13. Reads mapped to *N. viennensis* EN76 and *N. europaea* ATCC 19718 rRNA sequences were removed with SortMeRNA (v2.1) [32]. Next, the remaining reads were aligned to the genome of each strain using Bowtie2 (v2.4.4) [33], and the reads mapped to each gene were counted using HTSeq (v0.12.3) [34]. Expression values are presented as transcripts per kilobase million (TPM). The statistical analysis of differentially expressed genes was performed using the DESeq2 package in R.

### ROS scavenger experiment

We investigated the growth response of *N. viennensis* EN76 cells on floating filters in the presence of different ROS scavengers. Catalase (10 U mL-1) and varying concentrations of pyruvate (0 mM, 0.1 mM, and 1 mM) were used against H_2_O_2_-induced oxidative stress on the floating filter-grown cells compared to the control culture in liquid media supplemented with 0.1 mM pyruvate. Cells of *N. viennensis* EN76 with inoculum sizes of ∼10^7^ and 10^6^ cells were used for the experiment.

### Cultivation of AOA communities from agricultural soil

A composite bulk soil sample from an experimental agricultural station in Chungbuk National University Republic of Korea (127°27□18.5□E, 36°37□29.8□N) was used in this study. Sampling was done during the fallow period in March 2023. Six individual 30 cm subsoils were collected within a 10x10 m^2^ area from plots at the agricultural station. The subsoils were combined and transported to the laboratory and directly used for inoculation. The properties of the soil were as follows: loam texture (sand, 51%; silt, 33%; and clay, 16%); water content, 4.2%; pH, 6.0; total organic carbon, □0.1 g kg^-1^; total nitrogen, □0.01%; total ammonia, 6.0 mg kg^-1^; total phosphate, 174.8 mg kg^-1^; and cation exchange capacity, 12.1 cmol kg^-1^.

One gram (1 g) of soil was added into 10 mL AFM, vortex briefly, and serially diluted in 10-fold increments (i.e., 10^-1^ to 10^-4^). For comparison, one milliliter (1 mL) of each dilution was used as inoculum for floating filters and control culture in liquid media. For enrichment cultures, AFM was prepared with CaCO_3,_ and the final pH was 7.5. The soil suspension supplemented with the CaCO_3_ particles was used for inoculation. The protocol used to cultivate pure AOM strains on floating filters above was adopted for enriching soil AOM on floating filters. The petri dishes were placed in a sealed container and incubated in the dark at 30 °C without shaking. Nitrification activity was monitored by measuring NO_2_^-^ and NO_3_^-^ accumulation. NO_2_^-^ and NO_3_^-^ concentrations were quantified colorimetrically using the Griess and VCl_3_/Griess reagents, respectively [35]. For successive transfer of the enrichment cultures, cells grown on the filters were washed off, and 10% of the washed biomass was successively used as inoculum for new filters. For comparison, ca. 10% of the control culture in liquid media was repeatedly transferred for successive enrichment.

### 16S rRNA gene amplicon sequencing

For analysis of microbial communities enriched from agricultural soil, DNA extracted from the enrichment cultures and original inocula were used as templates for 16S rRNA gene amplicon sequencing analysis. A modified CTAB method [36] was employed in extracting high molecular weight genomic DNA from the enrichment cultures grown on floating filters and liquid media. In brief, biomass obtained from the cultures was treated with CTAB and sodium dodecyl sulfate (SDS) extraction buffer, incubated for 30 min at 65 °C with occasional mixing, and centrifuged at 8000 × g, 10 min, 25 °C. The supernatant obtained was repeatedly purified with an equal volume of chloroform/isoamyl alcohol (24:1). The extracted DNA was precipitated with 0.6 volume of 2-propanol, and the pelleted DNA was washed twice with 70% ethanol, allowed to air dry, and resuspended in TE buffer (10 mM Tris, pH 8, 1 mM EDTA). The extracted DNA concentrations were measured with a NanoDrop ND-1000 spectrophotometer (Thermo Fisher Scientific, Waltham, MA, United States), and the quality was assessed on a 1% (w/v) agarose gel.

The hypervariable V4-V5 region of the 16S rRNA gene was amplified with the primer pair 515F/926R [37] and sample indexing adapters (Nextera XT index kit). PCR amplifications were conducted via the following steps: 3 min heating step at 95 °C, followed by 25 cycles at 95 °C for 45 s (denaturation), 50 °C for 45 s (annealing), 72 °C for 90 s (extension), and 72 °C for 5 min (final extension). The PCR product was purified using the Labopass purification kit (Cosmo Genetech, South Korea), and the quantity obtained was measured with a NanoDrop ND-1000 spectrophotometer (Thermo Fisher Scientific, Waltham, MA, United States). The quality of the PCR product was assessed on a 1.5% (w/v) agarose gel. The library was sequenced using the Illumina MiSeq (2 × 300 bp) platform at Macrogen (Seoul, Republic of Korea). The QIIME (QIIME2-2023.2) pipeline with implemented tools for quality control (Cutadapt) [38], denoising and pair read merging (DADA2) [39], and de novo OTU clustering (VSEARCH) [40] was used to analyze the sequencing results. An annotated 16S rRNA reference dataset (SILVA release 132) [41] was used for taxonomic identification of the OTUs.

### Phylogenetic analysis

For phylogenetic analysis of the 16S rRNA gene sequence, representative nucleotide sequences of related taxa were obtained from the NCBI database. For phylogenetic analysis of sigma-70 protein sequences of *N. europaea* ATCC 19718, representative amino acid sequences of four diverse bacteria: *Escherichia coli*, *Caulobacter vibrioides*, *Bacillus subtilis*, *Pseudomonas aeruginosa* were also obtained from NCBI. The 16S rRNA gene sequences and the amino acid sequences of the sigma-70 protein were aligned separately with MAFFT [42]. Maximum-likelihood trees were inferred with IQ-TREE. The constructed trees were visualized using iTOL and used for annotation.

### Statistical Analysis

All statistical analyses were conducted using the R statistical software (v4.1.2) and R Studio (v2022.02.3). Microbial diversity analysis and data visualization were performed using the R packages phyloseq (v1.26.0) [43], vegan (V2.5-3) [44], and ggplot2 (v3.1.0) [45]. Processed amplicon sequence reads were imported using phyloseq [43]. Non-metric multidimensional scaling (NMDS) analysis, based on Bray-Curtis dissimilarity metrics, was used in Vegan [44] to compare the microbial communities between the samples. The ordination analysis patterns were statistically tested using permutational multivariate analysis of variance (PERMANOVA) and analysis of similarity (ANOSIM) with adonis2 and vegdist, respectively, which are part of the vegan packages in R [44]. Indicator species were identified using the indval function of labdsv package (v2.0-1) [46]. Samples from the same cluster were treated as a group for indicator species analysis.

## Results and Discussion

### Growth on floating filters

All the strains with varying cell densities were cultivated on floating filters, with the control culture in liquid media. The ammonia oxidation activity of *N. viennensis* EN76 cells with an inoculum size of ∼10^7^ cells decreased on floating filters compared with the control culture in liquid media (**Fig. 1A and B**). Furthermore, the ammonia oxidation activity of *N. viennensis* EN76 with an inoculum size of 10^5^ cells was significantly repressed on floating filters with less than < ca. 0.05 mM ammonia oxidized in 20 days (**Fig. 1A**). However, lower inoculum sizes (i.e.,∼10^6^ and 10^5^ cells) had no significant effect on the ammonia oxidation activity of the control culture grown in liquid media but almost proportionally increased the time required for complete oxidation of 1 mM NH_3_ (**Fig. 1B**). It is interesting that the difference in inoculum size only affects ammonia oxidation activity of *N. viennensis* EN76 cells grown on floating filters. On the contrary, □*N. chungbukensis*” MY2 cells showed no ammonia oxidation activity on floating filters compared to the control culture in liquid media (**Figs. 1C and D**). Nevertheless, the ammonia oxidation activity of *N. europaea* ATCC 19718 cells grown on floating filters was not notably different compared to the control culture in liquid media. Moreover, decreasing the inoculum sizes by 10-fold increments did not repress the ammonia oxidation activity of *N. europaea* ATCC 19718 cells grown on floating filters (**Figs. 1E and F**).

**Figure 1.**
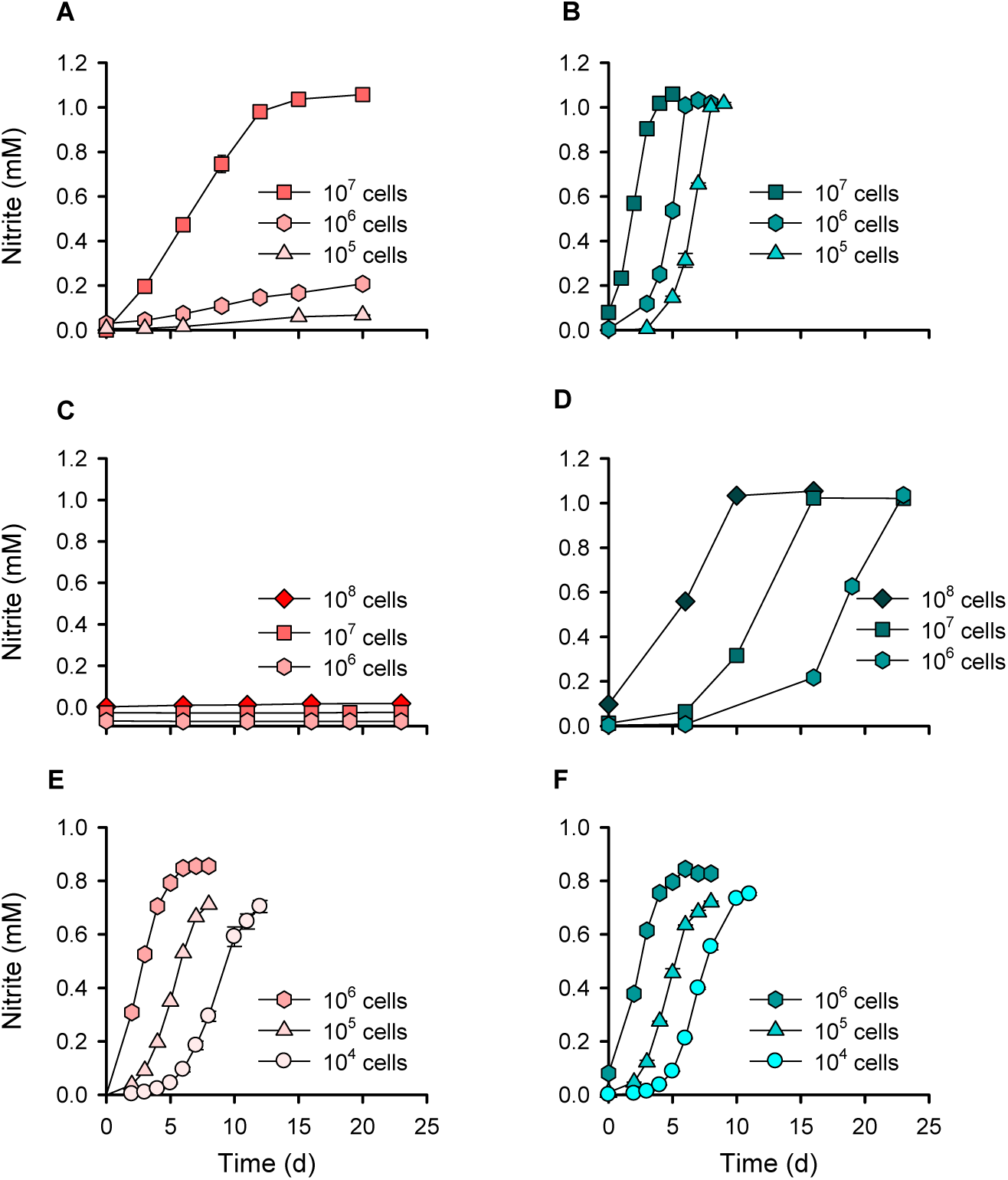
Comparison of ammonia oxidation activities of AOM grown on floating filters and the control culture in liquid media. The line graphs (**A**, **C**, **E**) and (**B**, **D**, **F**) represent floating filters and liquid cultures, respectively. *N. viennensis* EN76 (**A** and **B**), □*N. chungbukensis*” MY2 (**C** and **D**), and *N. europaea* ATCC 19718 (**E** and **F**) grown on floating filters and liquid media, respectively with varying inoculum size. All experiments were performed in triplicates. Data are presented as mean ± standard deviation (SD), and the error bars are hidden when they are smaller than the width of the symbols. To avoid overlapping symbols, the value was shifted by -0.03 and 0.02 in (**A**) for the experiment with 10^6^ cells and 10^5^ cells, respectively, and by -0.09, 0.06, or 0.02 in (**C**) for the 10^8^ cells, 10^7^ cells, and 10^6^ cells, respectively.

Considering no significant decrease in ammonia oxidization activity of *N. europaea* ATCC 19718 cells on floating filters, it can be inferred that any potential limitation in nutrient transport through the filters, which have ∼10^8^ pores/cm^2^, is likely insignificant, at least for *N. europaea* ATCC 19718. Since the number of pores on the filters is greater than the size of inoculated cells (<10^8^ cells/filter; filtered area approx. 11.3 cm^2^), it is improbable that the pores will become clogged by the inoculated cells. The effect of filter materials on the ammonia oxidation activity of *N. viennensis* EN76 cells with an inoculum size of ∼10^7^ cells was tested on other filters: polyvinylidene fluoride (PVDF), polytetrafluoroethylene (PTFE), and mixed cellulose esters (MCE). We observed comparable ammonia oxidation activities between the alternative filters (as mentioned above) and polycarbonate (PC) filters, indicating no significant inhibitory effect of filter materials (**Fig. S1**). Thus, the reason for the severe growth retardation in the lower inoculum sizes (∼10^6^ and 10^5^ cells) of *N. viennensis* EN76 cells on the surface of the filter is unclear. It is conceivable that an unknown cell density-dependent cooperative interaction and cellular responses [47] might be responsible. The volatilization of essential metabolic intermediates of ammonia oxidation, such as the gaseous NOx (HONO + NO + NO2) [48, 49], can also be serious for lower-density cultures on floating filters, which needs further investigation.

Using fluorescence microscopy, we observed microcolonies formation by *N. viennensis* EN76 with higher inoculum size (∼10^7^ cells) on the filters before and after oxidation of 1 mM NH_3_ (**Fig. 2A and B**), respectively. The size of microcolonies was further increased after oxidizing an additional 1 mM NH_3_ supplied into the AFM beneath the filters (**Fig. 2C**). However, the microcolony formation of *N. viennensis* EN76 with lower inoculum size (∼10^6^ cells) was poor compared with higher inoculum size (∼10^7^ cells) during the same incubation period (ca. 20 days) (**Fig. 2D**), corresponding with the observed decreased ammonia oxidation activity (**refer to Fig. 1A**). Furthermore, cellular protein quantification revealed that *N. viennensis* EN76 cells grown on floating filters produced less biomass than the control culture in liquid media (**Fig. S2**), implicating stress conditions for AOA grown on the filter.

**Figure 2.**
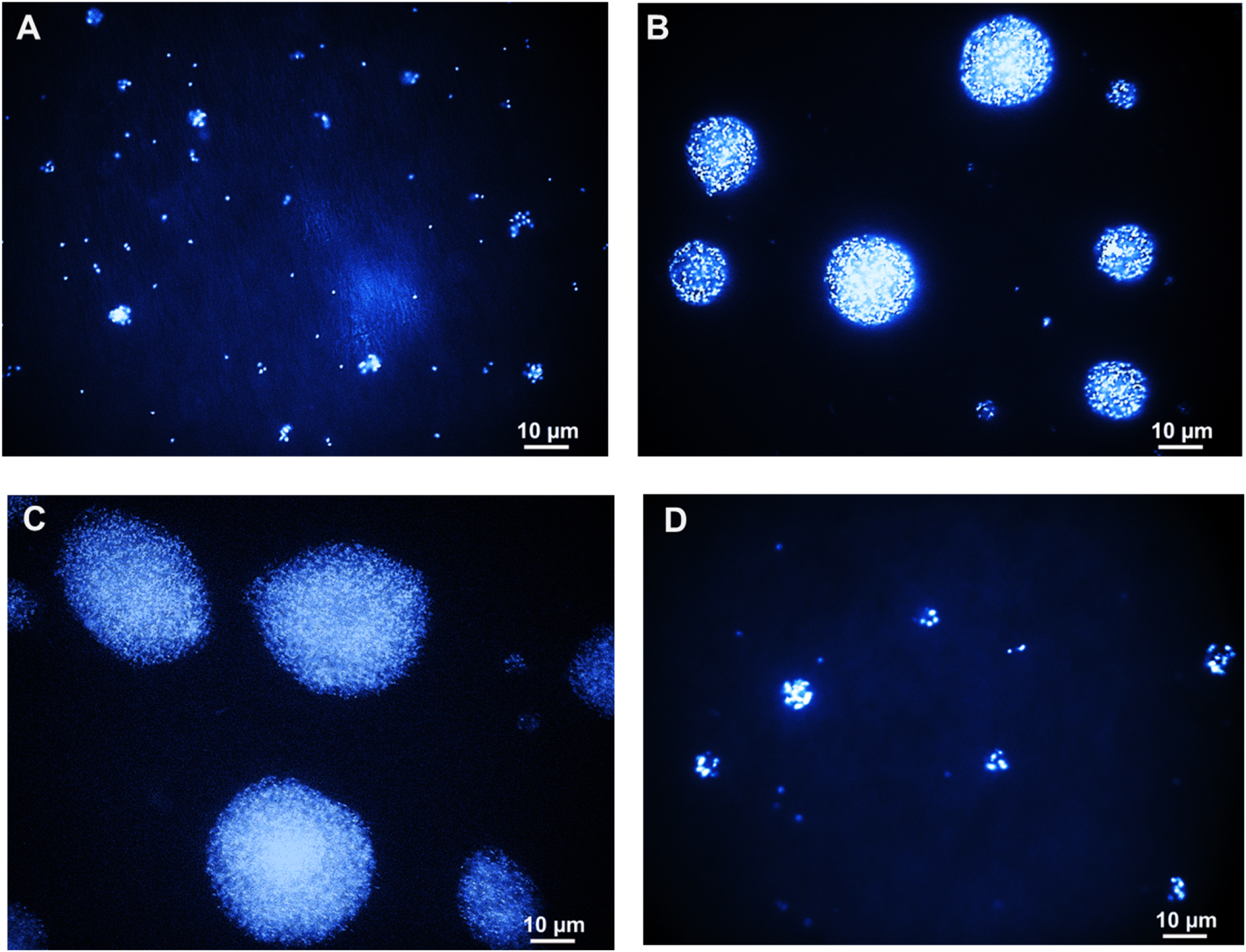
Fluorescent micrographs of *N. viennensis* EN76 microcolonies grown on floating filters. Micrographs of *N. viennensis* EN76 with an inoculum size of ∼10^7^ cells on polycarbonate filter (**A**) before and (**B**) after 1 mM ammonia oxidation (**refer to** Fig. 1A) and (**C**) microcolonies of *N. viennensis* EN76 after oxidation of an additional 1mM NH_3_. (**D**) Microcolonies of *N. viennensis* EN76 with an inoculum size of ∼10^6^ cells after 20 days of incubation (**refer to** Fig. 1A). The cells were stained with DAPI blue for 10 mins and dried at 37 °C on a glass slide.

### Effect of CaCO_3_ and filtration inoculation

Calcium carbonate (CaCO_3_) particles are often included in AOM cultures (liquid media) as a buffer against acid stress [50] and may increase the available surface that induces biofilm or microcolony formation in AOM [51, 52]. Owing to their deliquescent properties, CaCO_3_ particles surrounding the cells on the filters may act as a protective shield against dryness. Interestingly, we observed that inoculating lower inoculum size (∼10^6^ cells) of *N. viennensis* EN76 with CaCO_3_ particles on the filters improved the ammonia oxidation activity (**Fig. 3A**). In contrast, the growth of lower inoculum size (∼10^7^ cells) of □*N. chungbukensis*” MY2 on floating filters could not be recovered even with CaCO_3_ particles supplementation (**Fig. 3B**), indicating their inability to grow on the air-exposed filters or permanent damage of cells caused by the vacuum filtration process.

**Figure 3.**
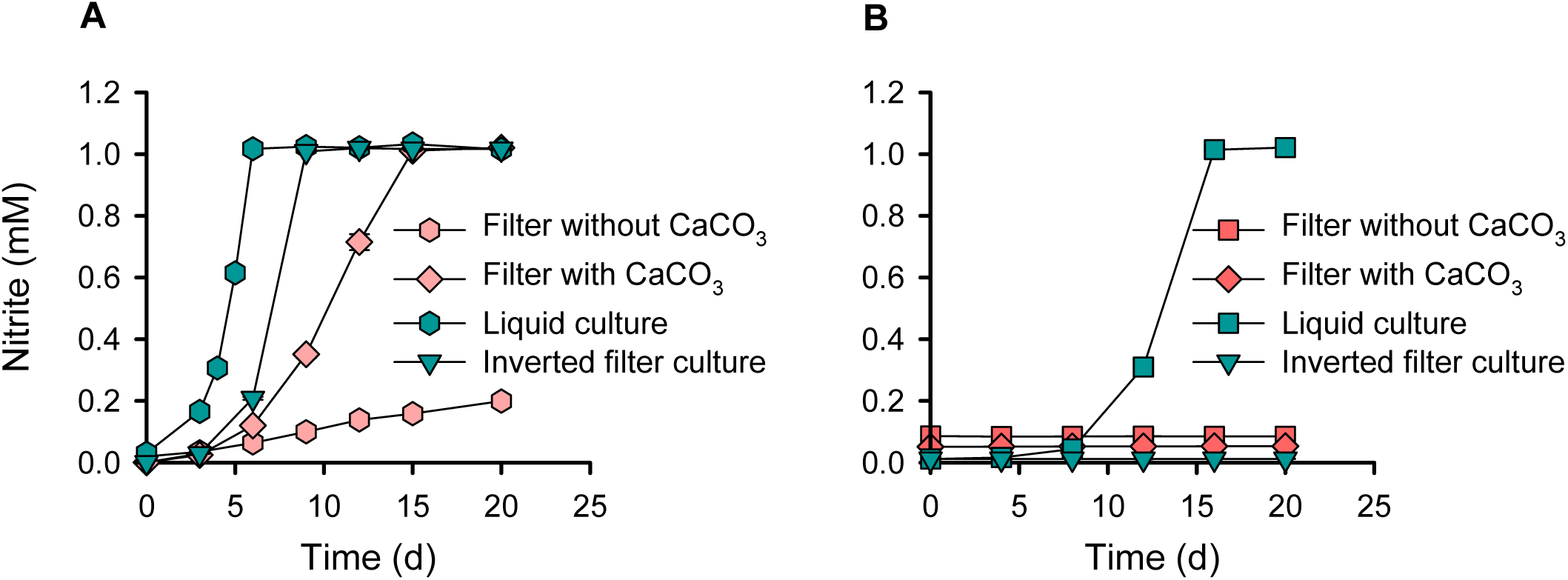
Ammonia oxidation activities of *N. viennensis* EN76 and □*N. chungbukensis*” MY2 cells under different growth conditions. *N. viennensis* EN76 (**A**) and □*N. chungbukensis*” MY2 (**B**) with inoculum sizes of ∼10^6^ cells and ∼10^7^ cells, respectively were used for the experiment. All experiments were performed in triplicates. Data are presented as mean□±□SD, and the error bars are hidden when they are smaller than the width of the symbols. To avoid overlapping symbols, the value was shifted by -0.02 in (**A**) for the filter without CaCO_3_ and by -0.09, 0.05, or 0.01 in (**B**) for the filter without CaCO_3_, filter with CaCO_3_, and inverted filter culture, respectively.

To assess the viability of □*N. chungbukensis*” MY2 cells after filtration, filters on which the cells were deposited were inverted, exposing cells directly to the AFM beneath the filter (i.e., inverted floating filter). However, □*N. chungbukensis*” MY2 cells showed no ammonia oxidation activity on inverted floating filters (**Fig. 3B**). On the contrary, the specific growth rate (*µ*_max_) of *N*. *viennensis* EN76 cells on inverted floating filters (0.59 ± 0.00 d^−1^) was comparable with the control culture in liquid media (0.63 ± 0.02 d^−1^) (**refer to Fig. 3A**). Nevertheless, the slight difference in the ammonia oxidation activity of *N. viennensis* EN76 cells on inverted floating filters and air-exposed floating filters (**refer to Fig. 3A**) further signifies a unique life strategy for AOA cells growing on air-exposed solid surfaces. Together, these results suggest that cells of □*N. chungbukensis*” MY2 of the *Nitrosopumilales* lost viability and are more vulnerable to physical stress damage than cells of *N. viennensis* EN76 of the *Nitrososphaerales*. This notion is supported by results from previous studies wherein □*N. chungbukensis*” MY2 and *Ca Nitrosopumilus maritimus* SCM1 of *Nitrosopumilales* lost their ammonia oxidation activities after the cells were concentrated by filtration or centrifugation [26, 53]. Accordingly, we observed ammonia oxidation activity from □*N. chungbukensis*” MY2 cells on floating filters when inoculated on filters using ambient gravitational force without vacuum suction (**Fig. S3**). Nonetheless, the observed ammonia oxidation activity of these cells was lower than the control culture in liquid media. This indicates that cells of □*N. chungbukensis*” MY2 may not be well adapted to the air-exposed solid surfaces, as it has been reported that AOA of *Nitrosopumilales* have limited capability to make extracellular polymeric substances (EPS), essential for biofilm or microcolony formation compared to AOA of *Nitrososphaerales* [54].

### Transcriptomic analysis of *N. viennensis* EN76

Considering the differences in ammonia oxidation activities (see above) of AOA cells grown on floating filters and the control culture in liquid media, we analyzed and compared the transcriptome of *N. viennensis* EN76 cells grown on floating filters and liquid media to better understand how AOA adapt to the air-exposed solid surfaces. Out of the 2,944 genes identified in the transcriptomes of *N. viennensis* EN76 (**Supplementary Table S1)**, 741 (25.17% of detected genes) exhibited significant differential expression with a log_2_FC > 1 and FDR < 0.05 (**Supplementary Table S2)**. Among the differentially expressed genes, 475 (64.1%) were significantly upregulated, and 266 (35.9%) were significantly downregulated in *N. viennensis* EN76 floating filter-grown cells. This differential gene expression indicates notable physiological differences between AOA cells grown as microcolonies on floating filters and the control culture in liquid media. Most differentially expressed genes in cells grown on floating filters could be functionally categorized in 1) Cell wall and extracellular polymeric substance (EPS) biosynthesis, 2) Inorganic ion transport and metabolism, 3) Posttranslational modification, protein turnover, and chaperones, 4) Carbohydrate transport and metabolism, 5) Signal transduction mechanisms. The ‘function unknown’ category contains the largest fraction of upregulated genes in floating filter-grown cells (**Supplementary Table S2 and Fig. S4**). Differentially expressed genes critically involved in the adaptation to growth on the filters were further analyzed.

#### Cell wall and EPS biosynthesis

The genes involved in EPS biosynthesis e.g., genes encoding glycosyl transferase, exported polysaccharide deacetylase, sialidase-neuraminidase family protein, methyltransferase, N-acetyltransferase, mannosyltransferase, sulfotransferase, and xylanase/chitin deacetylase [54], which often reside in clusters, were upregulated in floating filter-grown cells of *N. viennensis* EN76 (**Supplementary Table S3**). Some of these enzymes belong to the family of transporters (TC#4.D.1 and 4.D.2) that couple polysaccharides biosynthesis with translocation across the membrane. EPS production is essential for microcolony and biofilm development [55]. Thus, the functions of these enzymes may be critical for cell surface modification required for solid surface adaptation, thereby promoting survival by establishing interaction between cells and protecting cells from air-exposed environments. Besides EPS biosynthesis, genes encoding cell surface-associated proteins, such as hemolysin, pilin, and surface anchor family protein, were downregulated in floating filter-grown cells (**Supplementary Table S3**).

#### Oxidative stress responses/Inorganic nutrient homeostasis

Maintaining iron homeostasis is crucial in responding to oxidative stress caused by the Fenton reaction, where Fe(II) reacts with H_2_O_2_, generating a highly reactive and toxic hydroxyl radical (OH·) [56]. This reaction is primarily regulated by the mini-ferritins family of proteins known as Dps (DNA-binding protein from starved cells) [57, 58]. Dps proteins bind and store Fe(II), ferroxidizng it effectively using H_2_O_2_ as the oxidant while simultaneously detoxifying H_2_O_2_ [57, 59], thus preventing OH· production. Two of the genes encoding Dps proteins in *N. viennensis* EN76 (*dps3*; NVIE_RS09700 and *dps4*; NVIE_RS13730) were upregulated (> 6-fold) in floating filter-grown cells (**Supplementary Table S4**). Dps3 (NVIE_RS09700) in *N. viennensis* EN76 shares 68.16% identity with the Dps protein (WP_009989805.1) in *Saccharolobus solfataricus* (formerly referred to as *Sulfolobus solfataricus*). On the other hand, *N. viennensis* EN76 Dps4 (NVIE_RS13730) shares 24.00% and 40.76% identities with Dps protein from *Escherichia coli* (WP_000100800.1) and *Pseudomonas aeruginosa* PAO1 (NP_249653.1), respectively. Maaty et al. [60] reported high upregulation of the genes encoding Dps protein in *S. solfataricus* in response to H_2_O_2_-induced oxidative stress. Notably, the genes encoding Dps protein in *E. coli* and *P. aeruginosa* PAO1 are known to protect cells from H_2_O_2_-mediated oxidative stress [59, 61]. Therefore, the increased expression of these genes encoding Dps proteins in *N. viennensis* EN76 floating filter-grown cells indicates their involvement in iron homeostasis by decreasing the free iron that could catalyze the Fenton reaction. This process synchronously detoxifies H_2_O_2_, thereby preventing iron and H_2_O_2_-induced oxidative damage and promoting cell survival.

The gene encoding a rubrerythrin-like protein in *N. viennensis* EN76 (NVIE_RS11805) was upregulated (> 2-fold) in floating filter-grown cells (**Supplementary Table S4)**. Lumppio et al. [62] reported that the rubrerythrin protein in a sulfate-reducing anaerobic bacterium, *Nitratidesulfovibrio vulgaris* (formerly referred to as *Desulfovibrio vulgaris*), protects against exposure to air and H_2_O_2_-induced oxidative stress. Rubrerythrin protein can function as the terminal component of an NADH peroxidase, catalyzing the reduction of H_2_O_2_ to water [62]. The gene encoding the rubrerythrin-like protein in *N. viennensis* EN76 (NVIE_ RS11805) shares 38.71% and 30.65% identity with two rubrerythrin proteins (WP_010937330.1 and WP_010940353.1), respectively, in *N. vulgaris*. Previous studies [60, 63, 64] also showed that rubrerythrin proteins were upregulated in *Methanothermobacter thermautotrophicus*, *Porphyromonas gingivalis*, and *S. solfataricus* in response to H_2_O_2_-induced oxidative stress. In addition, genes encoding thioredoxin (*trxA1*; NVIE_RS13995 and *trxA2*; NVIE_RS14355), thioredoxin-like domain/NHL repeat-containing protein (*resA*; NVIE_RS00045), and peroxiredoxin-like protein (NVIE_RS05335) in *N. viennensis* EN76 were upregulated (> 2-fold) in floating filter-grown cells (**Supplementary Table S4)**. Collectively, upregulation of the genes encoding rubrerythrin-like, thioredoxin, and peroxiredoxin-like proteins in floating filter-grown cells of *N. viennensis* EN76 suggests their crucial role in protecting the cells against H_2_O_2_-induced oxidative stress. In contrast, the genes encoding alkyl hydroperoxide reductase (NVIE_RS05690 and NVIE_RS06670) and superoxide dismutase (NVIE_RS14475) in *N. viennensis* EN76 showed constitutive expression.

Copper is a vital trace element for aerobic organisms [65]. Bacteria possess a periplasmic CopC protein that binds copper and delivers it to the inner membrane CopD protein, which potentially transports copper into the cytoplasm [65, 66]. The two Cop proteins identified in the genome of *N. viennensis* EN76 are fusions of CopC and CopD domains, and their copper-binding site is highly conserved when aligned with other CopC and CopD representative sequences (**Fig. S5**). The genes (NVIE_RS06945 and NVIE_RS06955) encoding these proteins were upregulated (> 4-fold) in *N. viennensis* EN76 floating filter-grown cells (**Supplementary Table S4**). Intriguingly, *E. coli* cells exposed to high levels of copper showed less sensitivity to H_2_O_2_-induced DNA damage [67]. Recently, Guerra et al. [68] reported that cupric (Cu^2+^) ions occupy specific binding sites in Dps, thus exerting a significant rate-enhancing effect on ferroxidation reaction in aerobic microorganisms. This suggests that importing copper could serve as an effective mechanism for preventing or minimizing damage triggered by iron-induced OH· via classical Fenton reactions (as mentioned above). Similarly, a previous proteome study of three *Nitrosopumilus* strains (*Nitrosopumilus maritimus*, *N. adriaticus, N. piranensis*) revealed a relatively high abundance of Cop proteins in cells exposed to H_2_O_2_, implying a novel strategy for preventing oxidative damage through copper accumulation [69]. Therefore, the upregulation of the *copC/copD* genes in *N. viennensis* EN76 floating filter-grown cells may represent a means for copper acquisition and a unique approach to prevent oxidative cell damage, particularly on air-exposed floating filters. Together, our findings indicate that a distinct set of genes were expressed for H_2_O_2_-induced oxidative defense in soil AOA to thrive on air-exposed solid surfaces.

The genes encoding subunits A and B (*kdpA*; NVIE_RS12995 and *kdpB*; NVIE_RS12990) of the potassium translocating ATPase (Kdp) were upregulated (> 3- and 2-fold, respectively) in *N. viennensis* EN76 floating filter-grown cells, while the gene encoding subunit C (*kdpC*; NVIE_RS12985) was constitutively expressed (**Supplementary Table S4)**. The Kdp complex is a hybrid system for K^+^ transport that combines both the potassium (K^+^) superfamily transporters (KdpA) and the P-type ATPases (KdpB) [70, 71]. During the assembly of the Kdp complex, KdpC binds to KdpA to stabilize the complex, then K^+^ is transported from the KdpA subunit to the binding site in the KdpB subunit, where it is released to the cytosol [71]. The difference in K^+^ concentration across the plasma membrane plays a significant role in establishing the membrane potential, which is essential for regulating intracellular pH and generating the turgor pressure required for cell growth and division [72, 73]. The high affinity K^+^ uptake system, Kdp complex, is especially known to play critical roles in acid stress responses [74]. Nitrification causes acidification of media, which might be intensive on the floating filter. As observed above (**refer to Fig. 3A**), CaCO_3_ particles increased the ammonia oxidation activity of *N. viennensis* EN76 cells on floating filters. Thus, K^+^ homeostasis by the Kdp system is conceivably important for the acid adaptation of AOA on solid surfaces. It is notable that the representative acidophilic soil AOA, *Nitrosotalea* clade, as well as members of acid-adapted non-AOA soil *Nitrososphaerota*, *Gagatemarchaeaceae*, harbor the Kdp system [75, 76].

#### Ammonia oxidation

Most of the genes encoding different subunits of ammonia monooxygenase: *amoA*, *amoB*, *amoX*, *amoY*, *amoY*, *amoC1*, *amoC2*, *amoC4, amoC5,* and *amoC6* were constitutively expressed (**Supplementary Table S5**). Interestingly, the transcript of *amoC3* was highly upregulated (> 100-fold) in floating filter-grown cells of *N. viennensis* EN76 (**Supplementary Table S5**). It is important to note that *amoC3* and *amoC6* in *N. viennensis* EN76 are the core *amoC* COG in *Nitrososphaerales* [54]. Hodgskiss et al. [77] reported that *N. viennensis* EN76, like most other soil-dwelling AOA, encodes multiple homologs of the *amoC* gene and pinpointed *amoC6* as the primary homolog within the AMO enzyme complex. Depending on the environmental conditions, the various *amoC* homologs in the genome might provide different activity profiles to the AMO complex. Previous transcriptional studies showed that additional monocistronic *amoC* subunits found in some terrestrial AOB aid in maintaining AMO enzyme stability during stressful conditions [78–80]. Together, our findings suggest a pivotal role of the *amoC3* gene in the ammonia oxidation activity of *N. viennensis* EN76 grown on air-exposed solid surfaces.

The genome of *N. viennensis* EN76 encodes four of the type C two-domain multicopper oxidases (2dMCOs), with their copper-binding site (T1 center) overlaid onto the corresponding region in other type C 2dMCOs (**Fig. S6**). Notably, transcripts of MCOs were differentially expressed in *N. viennensis* EN76 cells grown on floating filters. MCO (NVIE_RS08635) and MCO (NVIE_RS12885) were significantly upregulated (>5- and 29-fold, respectively), while MCO (NVIE_RS00300) was downregulated in floating filter-grown cells. On the contrary, MCO (NVIE_RS09330) showed constitutive expression (**Supplementary Table S5**). In AOA, MCOs are predicted to participate in the archaeal HURM (hydroxylamine: ubiquinone redox) module, catalyzing the oxidation of hydroxylamine (the first product of ammonia oxidation) [81, 82]. Thus, it is conceivable that the MCOs (NVIE_RS08635 and NVIE_RS12885) might be involved in hydroxylamine oxidation in *N. viennensis* EN76 cells grown on air-exposed floating filters. *N. viennensis* EN76 cells grown on the filters seem to modify AMO and HURM systems to generate energy for surviving on solid surfaces.

#### Energy conservation

It has been suggested that electrons released into the electron transport chain (ETC) by hydroxylamine oxidation in AOA are transferred to the quinone/quinol pool (Q/QH_2_) via plastocyanin-like electron carriers [82, 83]. The floating filter-grown cells of *N. viennensis* EN76 showed significant upregulation (> 2-fold) of genes encoding the plastocyanin-like electron carriers and ferredoxins involved in electron transport (**Supplementary Table S6**). In addition, the genes encoding the *petC* (NVIE_RS09625) and *coxA2* (NVIE_RS00660) of Complex III and IV, respectively, were upregulated (> 4- and 15-fold, respectively) (**Supplementary Table S6)**. Electrons generated from ammonia oxidation are transferred to the bimetallic cytochrome a_3_/Cu_B_ active site in *coxA2*, where the reduction of dioxygen molecules to water occurs [84–86]. Furthermore, genes encoding the *atpC* (NVIE_RS11020) and *atpD* (NVIE_RS11000) of Complex V (ATP synthase), which are integral components of the catalytic site that utilizes proton motive force (PMF) for ATP synthesis, were downregulated in *N. viennensis* EN76 floating filter-grown cells (**Supplementary Table S6**). Meanwhile, other subunits of the ATP synthase showed constitutive expression. In the ETC, complex I (NADH-quinone oxidoreductase) is used for the reverse transport of electrons to reduce NAD^+^ in AOM [87–89]. Genes encoding six subunits (*nuoN*; NVIE_RS05590, *nuoL*; NVIE_RS05595, *nuoM*; NVIE_RS05600, *nuoJ*; NVIE_RS05610, *nuoI*; NVIE_RS05615, *nuoA*; NVIE_RS05640) of complex I were downregulated in *N. viennensis* EN76 floating filter-grown cells (**Supplementary Table S6**). The downregulation of these genes suggests reduced NADH production for anabolism. Thus, it is plausible that *N. viennensis* EN76 uses the activity of ETC to maintain PMF and sustain membrane potential for stress responses in cells grown on floating filters. The increased use of PMF has been reported in some bacteria under several starvation/stress conditions [90–92]. Differential expressions of genes involved in ammonium transport (**Supplementary Table S5**) and cell division (**Supplementary Table S7**) are described in **Supplementary Note 1.**

### H_2_O_2_ scavenger effect

The upregulation of genes involved in responding to H_2_O_2_-induced oxidative stress in *N. viennensis* EN76 cells grown on floating filters prompted us to investigate the effect of H_2_O_2_ scavengers on the floating filter culture. Typically, a concentration of 0.1 mM pyruvate is usually sufficient for scavenging H_2_O_2_ in liquid cultures of catalase-negative soil and marine AOA [17, 28]. To enhance H_2_O_2_ scavenging activity in the floating filter culture, we provided a higher concentration of pyruvate (1 mM) and catalase (10 U mL^-1^) into the media beneath the filter. Our results revealed that a higher concentration of pyruvate (1 mM pyruvate) and 10 U mL^-1^ catalase significantly increased the ammonia oxidation activity of *N. viennensis* EN76 cells grown on floating filters (**Fig. 4A and B**), indicating an improved H_2_O_2_ scavenging activity. Noticeably, the ammonia oxidation activity of *N. viennensis* EN76 cells with an inoculum size of ∼10^6^ cells increased significantly due to higher H_2_O_2_ scavenging activity (**Fig 4B**). Similarly, Bayer et al. [69] observed that higher initial cell density in *Nitrosopumilus* cultures could overcome H_2_O_2_-induced growth arrest in liquid media compared to cultures with lower cell density. This observation suggests that AOA cells exhibit a density-dependent cooperative defense against H_2_O_2_-induced oxidative stress. Together, the severe repression of ammonia oxidation activity of *N. viennensis* EN76 cells on floating filters (**refer to Fig. 1A**) may be attributed, in part, to H_2_O_2_-induced stress.

**Figure 4.**
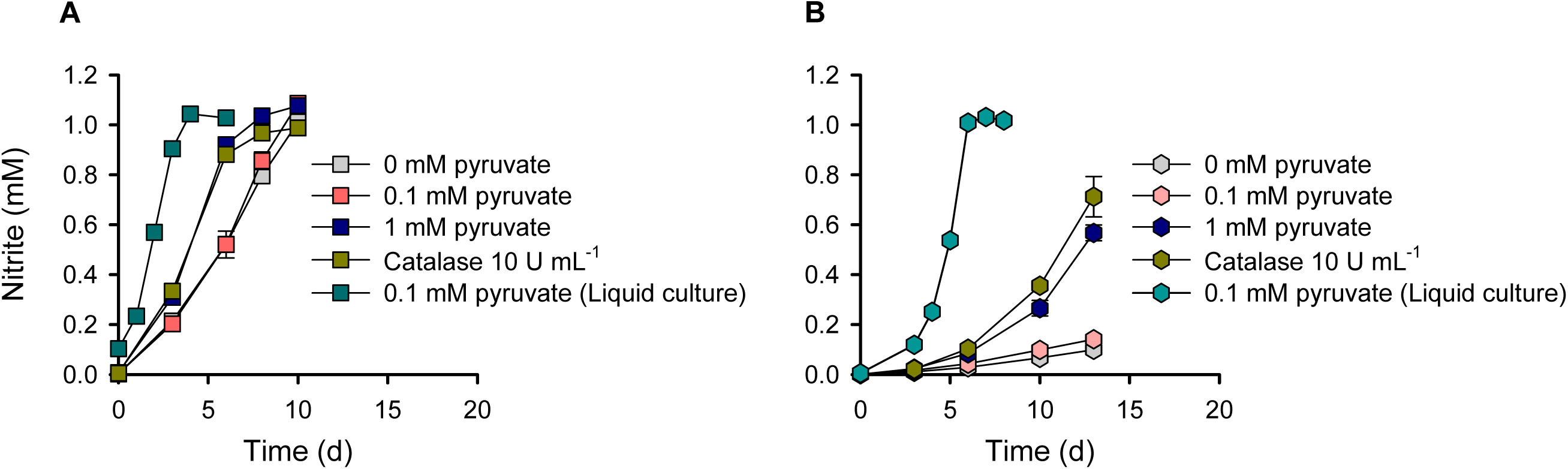
Effect of H_2_O_2_ scavengers on ammonia oxidation activity of *N. viennensis* EN76 cells grown on floating filters. Growth of *N. viennensis* EN76 cells with an inoculum size of (**A**) ∼10^7^ cells and (**B**) ∼10^6^ cells on floating filters provided with different concentrations of pyruvate and catalase (10 U mL^-1^), as compared to the standard pyruvate concentration used in liquid media. All experiments were performed in triplicates. Data are presented as mean□±□SD, and the error bars are hidden when they are smaller than the width of the symbols.

### Transcriptomic analysis of *N. europaea* ATCC 19718

We also compared the transcriptome of *N. europaea* ATCC 19718 cells grown on floating filters and the control culture in liquid media (**Supplementary Table S8**). Among the 2,619 genes identified in the transcriptome of *N. europaea* ATCC 19718, only 70 genes exhibited significant differential expression with a log_2_FC > 1 and FDR < 0.05. Prominently, genes encoding the sigma-70 (σ 70) family of the RNA polymerase (i.e., *fecI*), *fecR*-like genes, and the *fecA* (TBDT: TonB-dependent receptor) were the most abundant in upregulated genes in *N. europaea* ATCC 19718 floating filter-grown cells (**Supplementary Table S9**). Some genes involved in peptidoglycan synthesis and cell division were downregulated. The differential expression of a relatively small fraction of genes in *N. europaea* ATCC 19718 suggests that differences in the physiology of cells grown on floating filters and the control culture in liquid media are less significant when compared with *N. viennensis* EN76. This is consistent with the lack of notable differences in the ammonia oxidation activity of *N. europaea* ATCC 19718 between cells grown on floating filters and the control culture in liquid media (**refer to Figs. 1E** and **F**). Further description of the differentially expressed genes is provided and described in **Supplementary Note 2 and Fig. S7**.

### Cultivation of soil AOA on floating filter

The growth of AOA on floating filters provided valuable insights into their growth in air-exposed surfaces of soil aggregates. Thus, we adopted this technique to grow surface-adapted AOA from agricultural soil and compared it with those grown in liquid media. To focus on AOA communities, allylthiourea (ATU), an inhibitor of bacterial ammonia-oxidizers, was added to the growth media. Following the oxidation of about 0.7 mM NH_3_ in the first culture on floating filters and liquid media, 10% of the biomass was successively transferred to fresh filters and liquid media, respectively (**Figs. 5A and B**). Notably, there was no significant accumulation of NO_2_^-^ as an intermediate; instead, NO_3_^-^ was the product detected.

**Figure 5.**
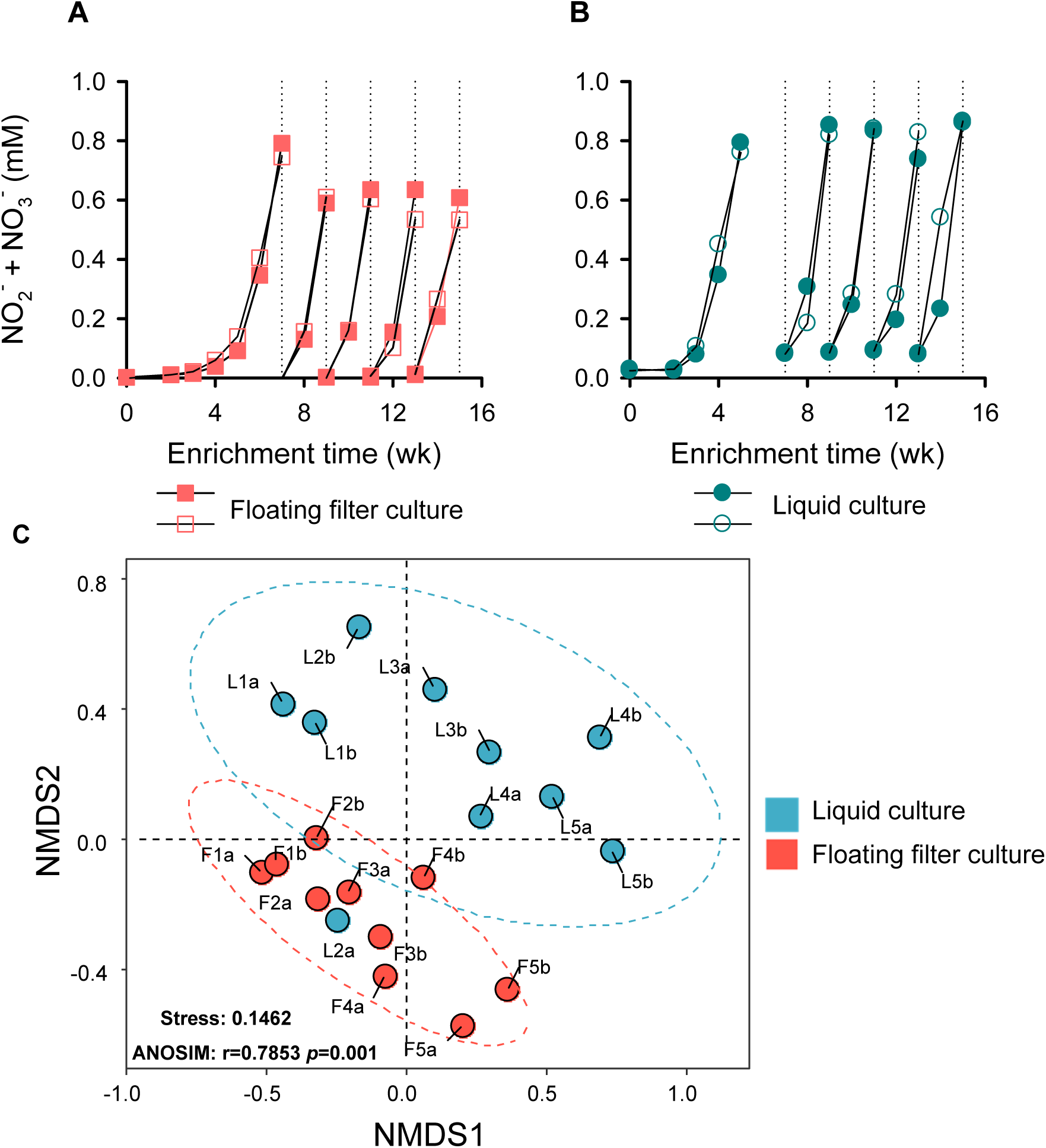
Cultivation of AOA from agricultural soil using the floating filter and liquid media cultivation technique. Experiments were conducted in the presence of ATU to inhibit the growth of AOB and comammox. Accumulation of NO_2_^-^ + NO_3_^-^ indicates ammonia oxidation activity in the cultures. The line graphs (**A** and **B**) represent the ammonia oxidation activities of two biological replicates of floating filters and liquid cultures, respectively, during successive culture transfers. A nonmetric multidimensional scaling (NMDS) analysis of the overall nitrifier communities enriched on floating filters and liquid media is shown in (**C**). OTU classification is based on 99% 16S rRNA gene amplicon sequence similarity. The letters and numbers indicate the following: L = liquid culture, F = floating filter culture, a & b = two biological replicates, 1 = first culture, 2 = 2nd culture, 3 = 3rd culture, 4 = 4th culture, and 5= 5th culture.

After four rounds of transfers, 16S rRNA gene amplicon sequencing was used to examine the nitrifier communities enriched on floating filters and liquid media. A nonmetric multidimensional scaling (NMDS) plot using the 16S rRNA gene operational taxonomic units (OTUs) with 99% similarity cutoff revealed significant variation (ANOSIM test, stress = 0.1462, R = 0.7853, P = 0.001) in nitrifier communities enriched on floating filters and liquid media during the transfers (**Fig. 5C**). The relative abundance of nitrifiers’ OTUs on floating filters was less than 20%, while the liquid media had a higher percentage of ca. 50% after four successive transfers. This difference implies that a significant amount of fixed carbon by AOM is used to cross-feed and support the growth of co-cultured heterotrophs [93] more on floating filters than liquid media (**Supplementary Table S10**). To identify the key OTU differentiating the nitrifier communities between floating filters and liquid cultures, an indicator species analysis was performed at the OTU level. Among the nitrifiers’ OTUs, only 16S_OTU4 was significantly more abundant on floating filters (IndVal: 0.6, *p* < 0.05) (**Table 1**). This OTU is closely related to members of the clade “*Ca*. Nitrosocosmicus” of the *Nitrososphaerales* lineage, with > 99.9% 16S rRNA sequence similarity (**Fig. 6A**). In contrast, 16S_OTUs (OTU30 and OTU52) within the *Nitrosopumilales* clades did not appear to thrive on floating filters (**Figs. 6B and C**), differing from their growth in liquid media (**Figs. 6D and E**). This is consistent with our previous observation of the weak growth of □*N. chungbukensis*” MY2 cells on floating filters (**refer to Figs. 1C and Fig. S3**). The 16S_OTU26 shows 99.19% 16S rRNA sequence similarity with *Nitrospira moscoviensis* BL23, indicating it might be the primary nitrite-oxidizing bacteria in floating filters and liquid cultures. The taxonomy of all nitrifiers’ OTUs is detailed in **Supplementary Table S11.**

**Figure 6.**
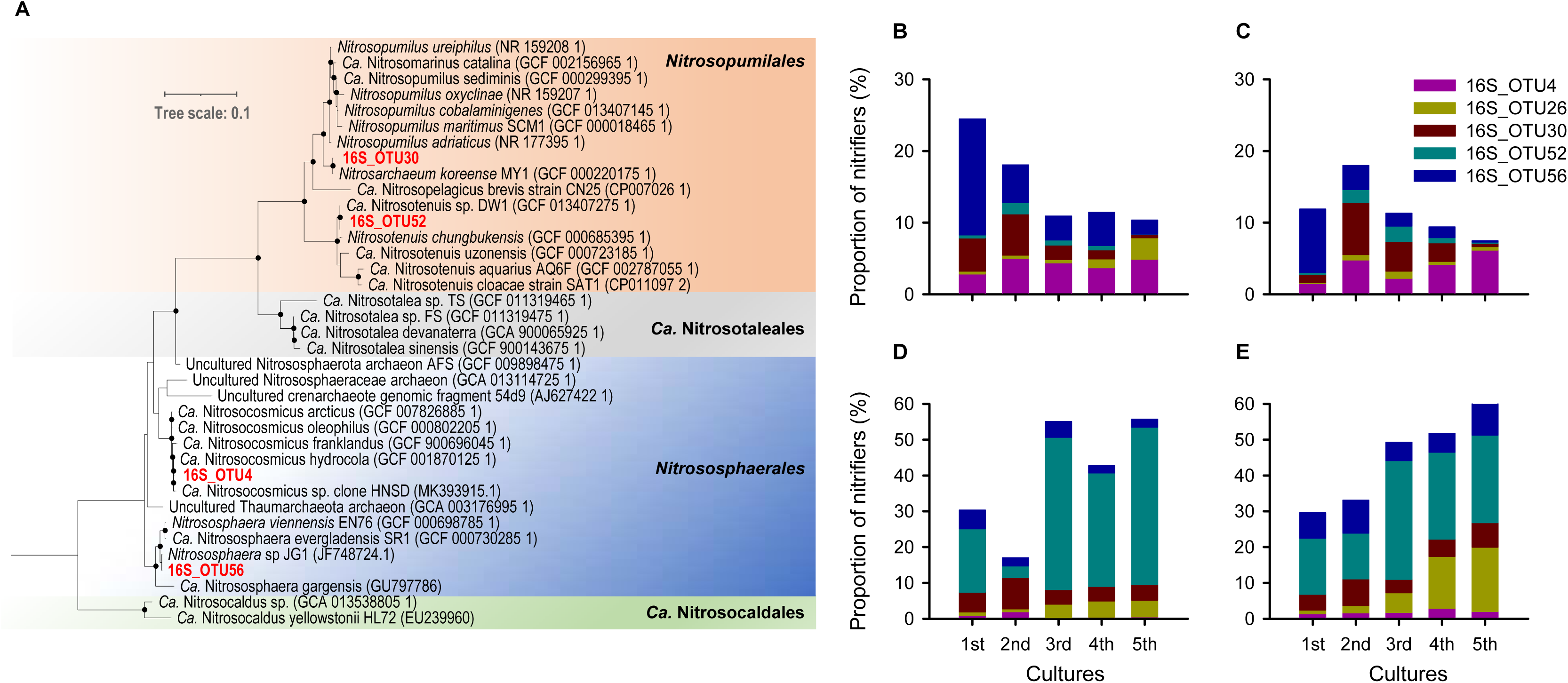
A phylogenetic tree of AOA_OTUs based on 16S rRNA gene sequences. Representative 16S rRNA sequences of AOA were selected from the National Center for Biotechnology Information databases. A maximum likelihood tree (**A**) was inferred with IQ-TREE (IQ-TREE options: -B 1000 -m LG+F+R5) using aligned sequences rooted at the mid-point. Bootstrap values ≥ 70% based on 1,000 replications are indicated. The scale bar represents a 0.1 change per nucleotide position. OTUs obtained in this work are red-coloured. Charts (**B**, **C**) and (**D**, **E**) represent the composition of nitrifiers of two biological replicates of floating filters and liquid cultures, respectively. The 16S_OTU26 is affiliated with the genus *Nitrospira*. Details of all nitrifiers’ OTUs taxonomy are provided in **Supplementary Table S11**.

**Table 1.**
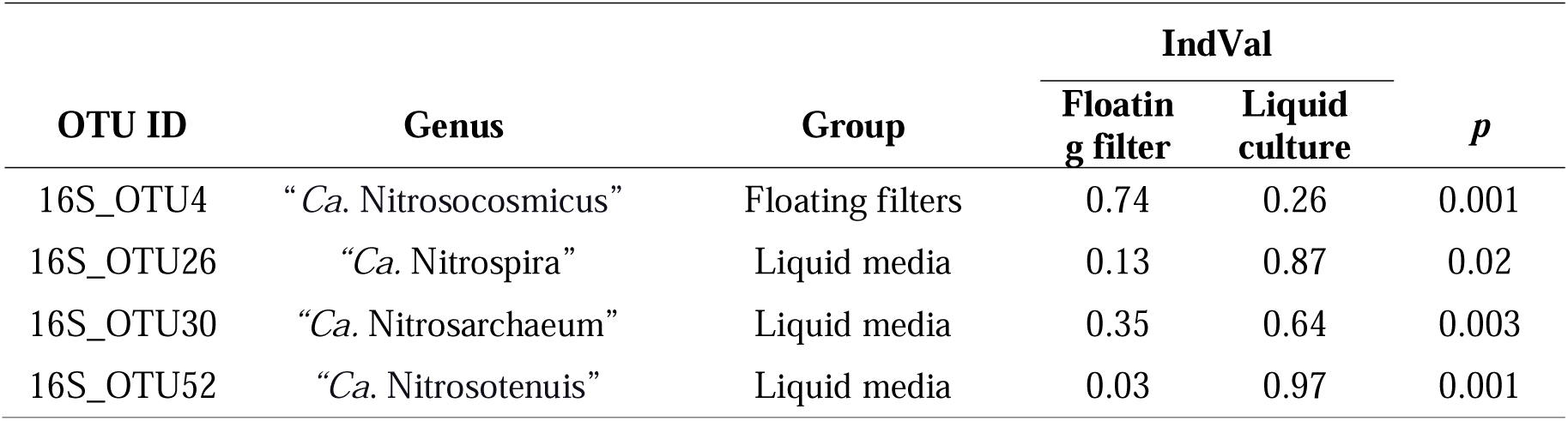
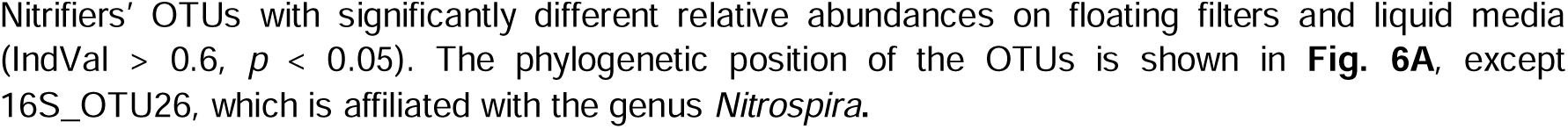
Indicator species analysis showing the nitrifiers OTUs on floating filters and liquid media.

| OTU ID | Genus | Group | IndVal |  | *p* |
| --- | --- | --- | --- | --- | --- |
|  |  |  | Floating filter | Liquid culture |  |
| 16S_OTU4 | “*Ca. Nitrosocosmicus*” | Floating filters | 0.74 | 0.26 | 0.001 |
| 16S_OTU26 | “*Ca. Nitrospira*” | Liquid media | 0.13 | 0.87 | 0.02 |
| 16S_OTU30 | “*Ca. Nitrosarchaeum*” | Liquid media | 0.35 | 0.64 | 0.003 |
| 16S_OTU52 | “*Ca. Nitrosotenuis*” | Liquid media | 0.03 | 0.97 | 0.001 |

Unlike most other catalase-negative AOA, members of “*Ca*. Nitrosocosmicus” harbor a gene encoding manganese catalase [16, 18], which may confer their resistance to H_2_O_2_-induced oxidative stress while growing on floating filters. In addition, the Kdp system in the “*Ca*. Nitrosocosmicus” clade [76, 94] and their resilience to high saline conditions [95] further support their potential adaptation to soil surfaces. Together, our results indicate that the “*Ca*. Nitrosocosmicus” clade might be adapted to air-exposed soil surfaces, and the floating filter cultivation technique can be exploited to obtain novel soil-dwelling nitrifiers.

### Conclusion

In this study, we demonstrated the ability of a soil ammonia-oxidizing archaeon, *N. viennensis* EN76, to thrive on air-exposed solid surfaces and compared its growth with that of an ammonia-oxidizing bacterium, *N. europaea* ATCC 19718. The physiological and transcriptional responses of these microorganisms highlight that the physiology of *N. viennensis* EN76 grown on solid surfaces (floating filter-grown cells) is notably different from the control culture in liquid media. *N. viennensis* EN76 exhibited significant upregulation of genes involved in the cell wall and EPS biosynthesis, H_2_O_2_-induced oxidative stress response, and ammonia oxidation when cultivated on floating filters. These adaptations are likely crucial for its survival and functionality in air-exposed solid surfaces.

Ecologically, this study underscores the importance of surface-attached growth for soil AOA. Given that soil microorganisms predominantly exist as microcolonies or biofilms within pores of soil aggregates directly exposed to the atmosphere, the ability of AOA to thrive in such conditions is critical for their roles in nitrogen cycling. Thus, the floating filter cultivation approach has proven to be a valuable tool for studying the ecophysiology of soil AOA, providing insights that are more representative of their natural habitats than traditional liquid culture methods. Furthermore, the observed distinct soil AOA communities dominated by catalase-containing AOA enriched on filters compared to liquid culture suggest that the floating filter cultivation technique can indeed potentially cultivate soil surface-adapted nitrifiers. This approach presents an exciting opportunity for further exploration of the function, activity, and diversity of AOA communities in various soil environments. Overall, this study sheds light on the adaptive mechanisms governing AOA growth on air-exposed solid surfaces and its ecological relevance, paving the way for more accurate and ecologically valid investigations of the soil nitrification process.

## Data Availability

The whole-transcriptome data and 16S rRNA gene amplicon sequencing data generated in this study have been deposited in the NCBI BioProject database under the accession project number (PRJNA1131856) and (PRJNA1131940), respectively.

## Acknowledgments

This work was supported by the NRF (National Research Foundation of Korea) grant funded by the Korean government (Ministry of Science and ICT) (2021R1A2C3004015) and Basic Science Research Program through NRF funded by the Ministry of Education (2020R1A6A1A06046235). J-HG was supported by the NRF grant funded by the Korean government (Ministry of Science and ICT) (RS-2023-00213601). M-YJ was supported by the NRF grant funded by the Korean government (Ministry of Science and ICT) (2021R1C1C1008303 and 2022R1A4A503144711).

