## Supplementary information for "Growth of soil ammonia-oxidizing archaea on air-exposed solid surface"

*Sung-Keun Rhee

**The file includes:**

Supplementary Notes 1 and 2

Supplementary Figures S1 to S7

Legends for Supplementary Tables S1 to S11

Supplementary References.

**Supplementary Note 1**

#### **Transcriptome analysis of N. viennensis EN76.**

***Ammonium transport and Cell division:*** Two of the ammonium transporters (*amt1*; NVIE_RS01185, and *amt3*; NVIE_RS11415) were downregulated in *N.* *viennensis* EN76 floating filter-grown cells, while *amt2* (NVIE_RS10630) showed constitutive expression (**Supplementary Table S5**). These results correspond with the lower growth rates and biomass production of *N.* *viennensis* EN76 cells grown on floating filters compared to the control culture in liquid media. The Cdv system is the primary cell division system in archaea, as evidenced in a marine ammonia-oxidizing archaeon, *Nitrosopumilus* *maritimus* SCM1 [1]. We observed that the putative archaeal cell division genes (*cdvB*; NVIE_RS02340 and *cdvC*; NVIE_025520) were downregulated in *N.* *viennensis* EN76 floating filter-grown cells (**Supplementary Table S7**). In contrast, other genes of the Cdv system such as *cdvA* (NVIE_RS00655), other *cdvB* homologs (*snf7*; NVIE_RS09745, *snf7*; NVIE_RS04040, *snf7*; NVIE_RS09110, and *snf7*; NVIE RS12195), and *ftsZ* (NVIE_RS05305) were constitutively expressed (**Supplementary Table S7**). Caspi et al. [2] reported that *cdvB* and *cdvC* are crucial for the endosomal sorting complex required for transport (ESCRT) pathway, which is involved in various cell division processes. Therefore, downregulation of the *cdvC*, which encodes an ATPase essential for the turnover of ESCRT membrane-abscission polymers [2], may lead to reduced turnover of these polymers, consequently affecting the regulation of controlled division of cells grown on floating filters.

**Supplementary Note 2**

### **Transcriptomic analysis of *N. europaea* ATCC 19718**

**T**he key genes involved in ammonia oxidation activity and energy conservation showed constitutive expression in *N. europaea* ATCC 19718 (**Supplementary Table S8**). Among the 70 genes that exhibited significant differential expression **(**log_2_FC > 1 and FDR < 0.05), 53 were upregulated, and 17 were downregulated in *N. europaea* ATCC 19718 floating filter-grown cells (**Supplementary Table S9**).

Among the sigma-70 (σ 70) families, group IV, also known as the extracytoplasmic function sigma factor (σ ECF) subfamily, is the largest and most diverse subfamily [3, 5]. A phylogenetic analysis of the nine upregulated σ 70 sequences in cells of *N. europaea* ATCC 19718 grown on floating filters (**Fig. S7**) revealed their clustering within the σ ECF subfamily. Cell-surface signaling, a signaling cascade that starts at the outer membrane and ends in the cytoplasm, was first described for the FecI/FecR pair in iron acquisition in *E. coli* [6]. The regulatory system was expanded to cell envelop-related processes such as biofilm formation and iron metabolism [6–9] This finding aligns with our observation in floating filter-grown cells of *N. viennensis* EN76, where the genes for EPS biosynthesis and iron homeostasis were upregulated. Together, the extracytoplasmic stress response mediated by the σ ECF subfamily in *N. europaea* ATCC 19718 cells grown on floating filters might be successfully used to sustain ammonia oxidation activity and growth of *N*. *europaea* ATCC 19718, as observed in Fig. 1E.

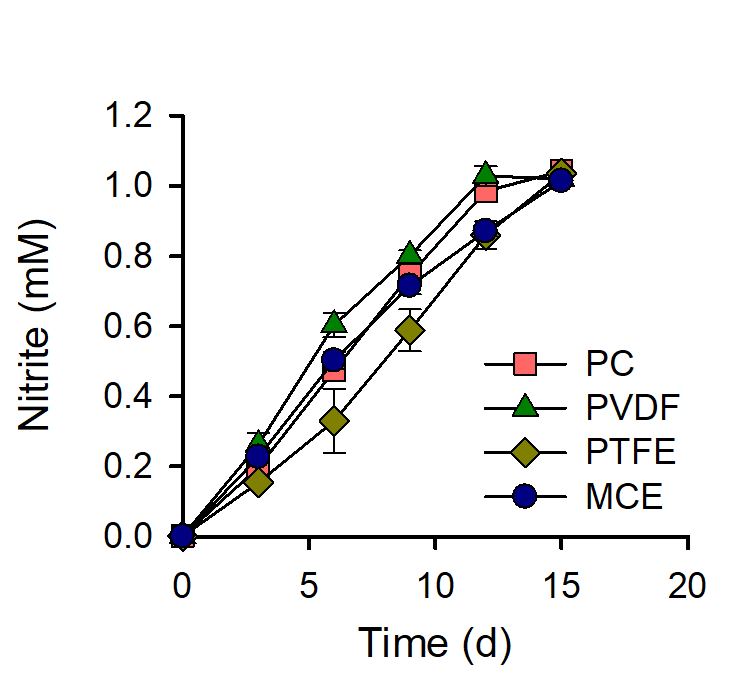

**Figure S1.** Ammonia oxidation activity of *N.* *viennensis* EN76 cells grown on different filters. The filters used include PC: polycarbonate, PVDF: polyvinylidene fluoride, PTFE: polytetrafluoroethylene, and MCE: mixed cellulose esters. An inoculum size of 10^7^ cells was used for the experiment. All experiments were performed in triplicates. Data are presented as mean ± SD, and the error bars are hidden when they are smaller than the width of the symbols.

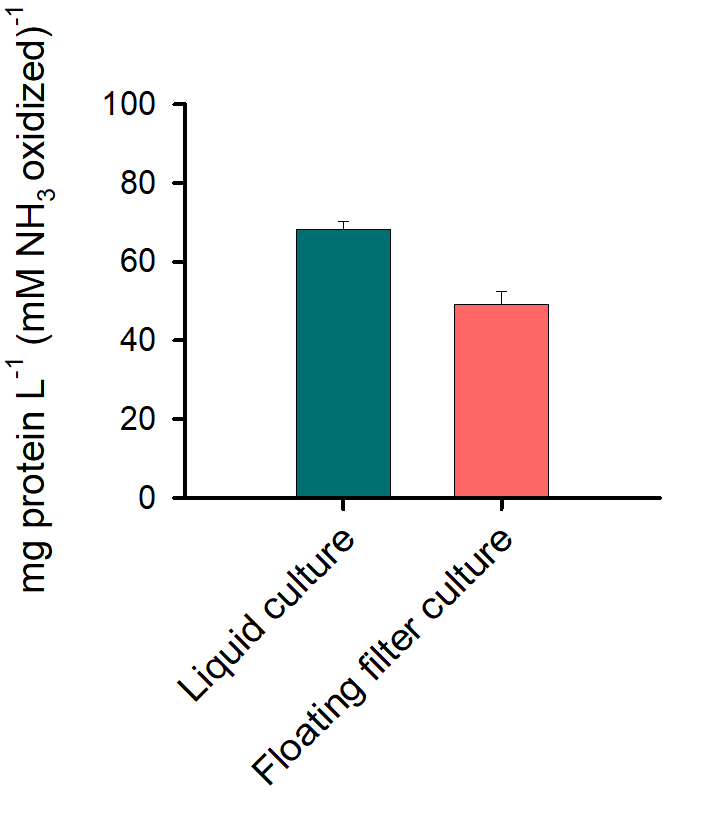

**Figure S2.** Growth yield of *N. viennensis* EN76 cells grown on floating filters and liquid media. The yield was calculated as total cellular proteins produced after complete oxidation of 1 mM ammonia. All experiments were performed in triplicates. Data are presented as mean ± SD.

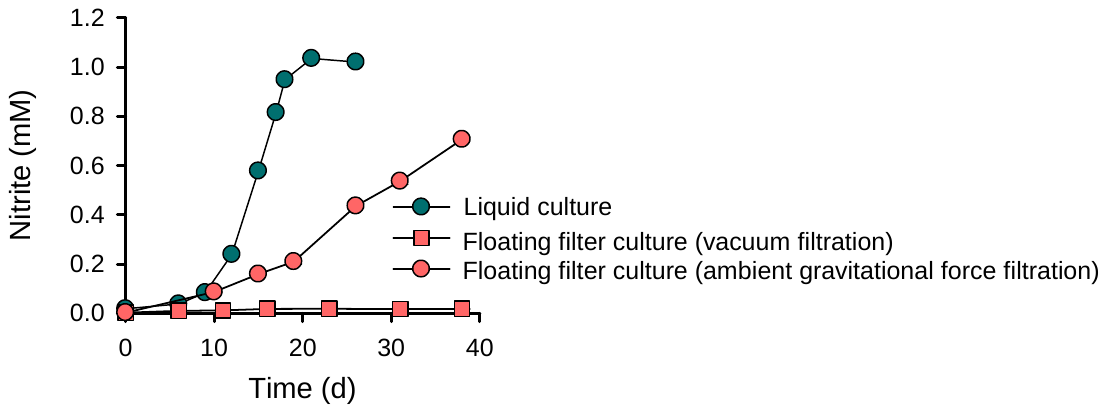

**Figure S3.** Ammonia oxidation activity of ‟*N. chungbukensis*” MY2 cells inoculated on floating filters using vacuum and ambient gravitational force filtration. Ammonia oxidation in the control culture grown in liquid media was used for comparison. All experiments were performed in triplicates. Data are presented as mean ± SD, and the error bars are hidden when they are smaller than the width of the symbols.

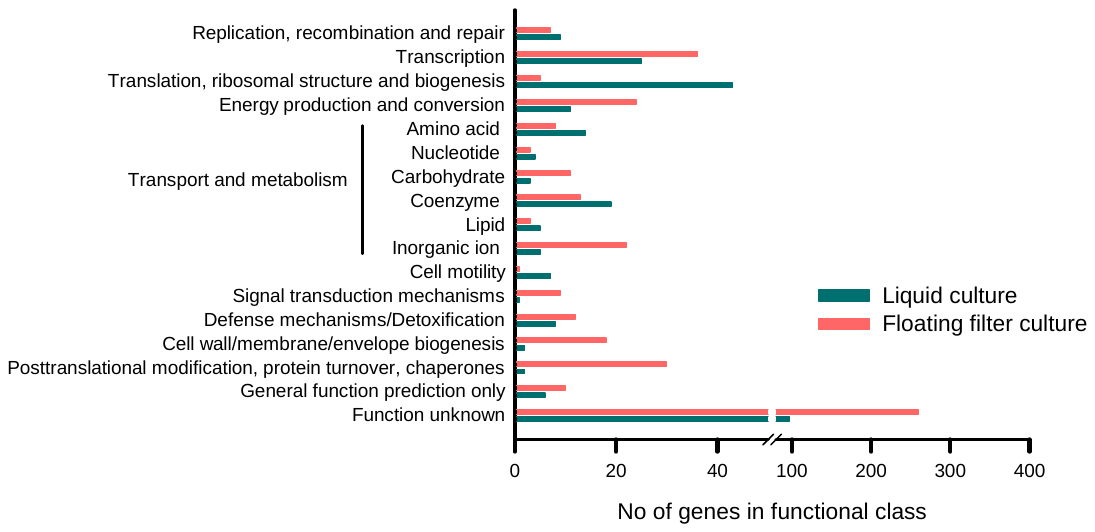

**Figure S4.** Comparative gene expression pattern of *N.* *viennensis* EN76 cells grown on floating filters and liquid media. Each bar in the functional class shows the number of upregulated genes. Genes are assumed to be differentially expressed when the log_2_FC > 1 and FDR < 0.05. Transcriptome experiments were performed in triplicates.

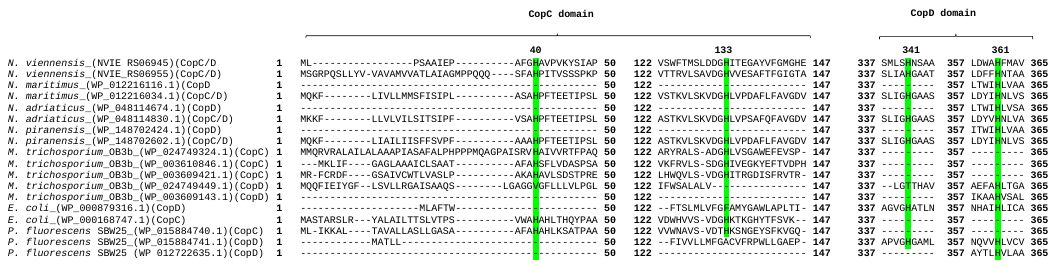

**Figure S5.** Alignment of the amino acid sequences of the Cop proteins (CopC*/*CopD NVIE_RS06945 and NVIE_RS06955) from *N. viennensis* EN76 with other Cop proteins. CopC and CopD representative sequences from *Nitrosopumilus* *maritimus* (WP_012216116.1 and WP_012216034.1), *N. adriaticus* (WP_048114674.1 and WP_048114830.1), *N. piranensis* (WP_148702424.1 and WP_148702602.1), *Methylosinus trichosporium* OB3b (WP_024749324.1, WP_003610846.1, WP_003609421.1, WP_024749449.1 and WP_003609143.1), *Escherichia coli* (WP_000879316.1 and WP_000168747.1), and *Pseudomonas* *fluorescens* SBW25 (WP_015884740.1, WP_015884741.1 and WP_012722635.1) were included. The numbers above and on the side of each sequence fragment indicate the residue position. The green colour in the alignments indicates the consensus positions of the copper-binding site.

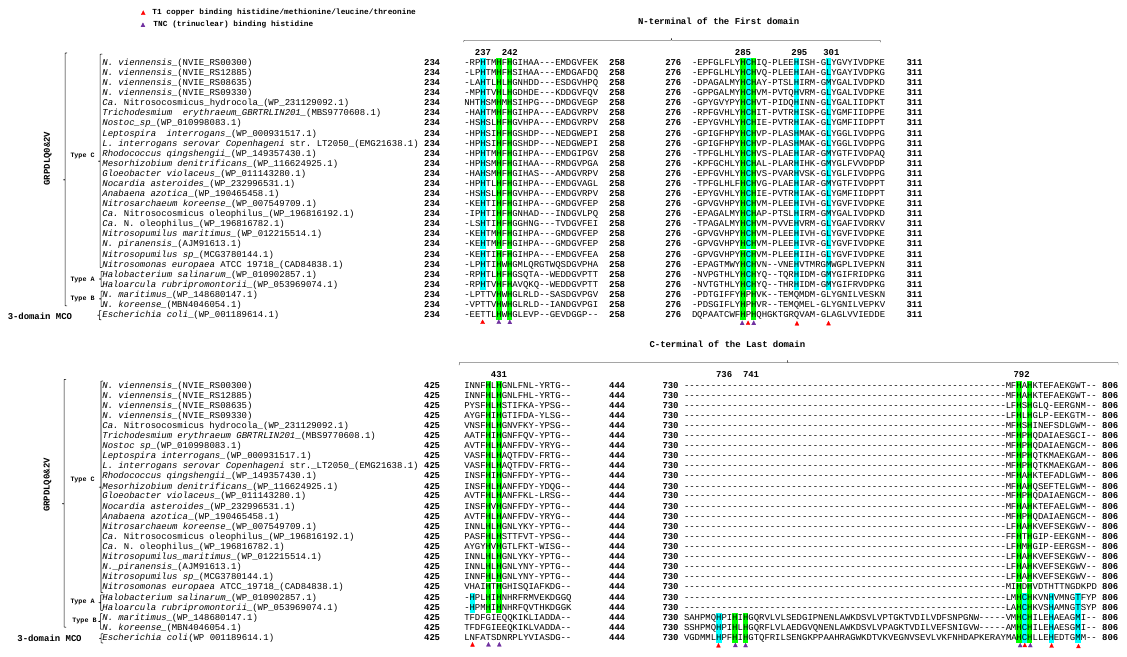

**Figure S6.** Alignment of amino acid sequence of multicopper oxidases (MCOs) showing the copper-binding sites of two-domain and three-domain. Sequences of two-domain MCOs (2dMCOs) are clustered into type A, B, and C. Two sequence fragments, from the N-terminal of the first and C-terminal of the last domains of each MCO, are indicated. The numbers above and on the side of each sequence fragment indicate the residue position. The turquoise and green colours in the alignments indicate the consensus positions of the copper-binding residues and the trinuclear histidine binding sites, respectively. Residues with the red-colored triangles below them are of T1 copper sites. Residues with purple-colored triangles below them are trinuclear binding histidine. The sequence ID (NCBI accession number) and the origin of each sequence are listed. This figure was modified from Figure 1 in [10].

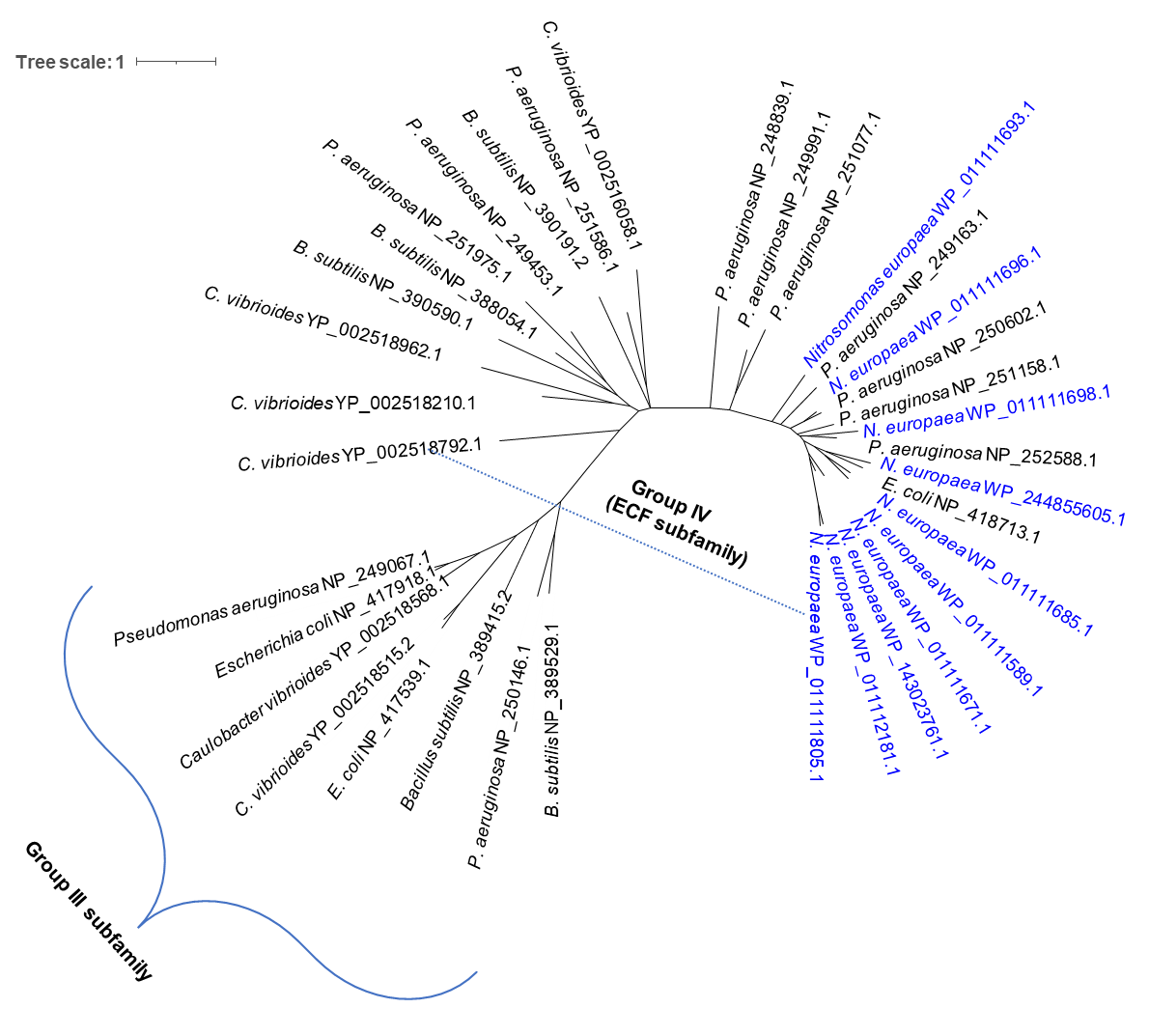

**Figure S7.** Maximum-likelihood phylogenetic analysis of the nine upregulated sigma-70 protein sequences from *N. europaea* ATCC 19718 floating filter-grown cells. Sigma-70 protein sequences in some other bacterial strains were selected from the National Center for Biotechnology Information databases. The unrooted tree was constructed with IQ-TREE (IQ-TREE options: -B 1000 -m LG+F+R5) using aligned sigma-70 sequences (details in *Materials and Methods*). The scale bar represents 1 change per amino acid position. The sequence ID (NCBI accession number) and the origin of each sequence are given for all strains. All the sigma-70 protein sequences from *N. europaea* ATCC 19718 are phylogenetically related to the group IV (sigma ECF) subfamily.

**Supplementary Tables**

**Supplementary Table S1:** List of all the genes expressed in *N. viennensis* EN76 grown on floating filters and liquid media.

**Supplementary Table S2:** Genes considered significantly upregulated or downregulated in *N. viennensis* EN76 grown on floating filters and liquid media. The threshold values of differentially expressed genes were set as log_2_FC > 1 and FDR < 0.05.

**Supplementary Table S3:** Differentially expressed genes involved in cell surface modification, EPS production, motility, and adhesion in *N. viennensis* EN76 grown on floating filters and liquid media. The threshold values of differentially expressed genes were set as log_2_FC > 1 and FDR < 0.05.

**Supplementary Table S4:** Differentially expressed genes involved in oxidative stress and inorganic nutrient homeostasis in *N. viennensis* EN76 grown on floating filters and liquid media. The threshold values of differentially expressed genes were set as log_2_FC > 1 and FDR < 0.05.

**Supplementary Table S5:** Differentially expressed genes involved in ammonia oxidation in *N. viennensis* EN76 grown on floating filters and liquid media. The threshold values of differentially expressed genes were set as log_2_FC > 1 and FDR < 0.05.

**Supplementary Table S6:** Differentially expressed genes involved in the electron transport chain in *N. viennensis* EN76 grown on floating filters and liquid media. The threshold values of differentially expressed genes were set as log_2_FC > 1 and FDR < 0.05.

**Supplementary Table S7:** Differentially expressed genes involved in cell division in *N. viennensis* EN76 grown on floating filters and liquid media. The threshold values of differentially expressed genes were set as log_2_FC > 1 and FDR < 0.05.

**Supplementary Table S8:** List of all the genes expressed in *N. europaea* ATCC 19718 grown on floating filters and liquid media.

**Supplementary Table S9:** Genes significantly upregulated or downregulated in *N. europaea* ATCC 19718 grown on floating filters and liquid media. The threshold values of differentially expressed genes were set as log_2_FC > 1 and FDR < 0.05.

**Supplementary Table S10**: Relative abundance of 16S rRNA OTUs of prokaryotes enriched on floating filters and liquid media.

**Supplementary Table S11**: Relative abundance of 16S rRNA OTUs of nitrifiers enriched on floating filters and liquid media.
